# Refinement of the “*Candidatus* Accumulibacter” Genus Based on a Metagenomic Analysis of Biological Nutrient Removal (BNR) Pilot-Scale Plants Operated with Reduced Aeration

**DOI:** 10.1101/2023.11.20.567894

**Authors:** Rachel D. Stewart, Kevin S. Myers, Carly Amstadt, Matt Seib, Katherine D. McMahon, Daniel R. Noguera

## Abstract

Members of the “*Candidatus* Accumulibacter” genus are widely studied as key polyphosphate-accumulating organisms (PAOs) in biological nutrient removal (BNR) facilities performing enhanced biological phosphorus removal (EBPR). This diverse lineage includes 18 “*Ca*. Accumulibacter” species, which have been proposed based on the phylogenetic divergence of the polyphosphate kinase 1 (*ppk1*) gene and genome-scale comparisons of metagenome-assembled genomes (MAGs). Phylogenetic classification based on the 16S rRNA genetic marker has been difficult to attain because most “*Ca*. Accumulibacter” MAGs are incomplete and often do not include the rRNA operon. Here, we investigate the “*Ca*. Accumulibacter” diversity in pilot-scale treatment trains performing BNR under low dissolved oxygen (DO) conditions using genome-resolved metagenomics. Using long-read sequencing, we recovered medium and high-quality MAGs for 5 of the 18 “*Ca*. Accumulibacter” species, all with rRNA operons assembled, which allowed a reassessment of the 16S rRNA-based phylogeny of this genus and an analysis of phylogeny based on the 23S rRNA gene. In addition, we recovered a cluster of MAGs that based on 16S rRNA, 23S rRNA, *ppk1*, and genome-scale phylogenetic analyses do not belong to any of the currently recognized “*Ca*. Accumulibacter” species for which we propose the new species designation “*Ca*. Accumulibacter jenkinsii” sp. nov. Relative abundance evaluations of the genus across all pilot plant operations revealed that regardless of the operational mode, “*Ca*. A. necessarius” and “*Ca*. A. propinquus” accounted for more than 40% of the “*Ca*. Accumulibacter” community, whereas the newly proposed “*Ca*. A. jenkinsii” accounted for about 5% of the “*Ca*. Accumulibacter” community.

**IMPORTANCE:** One of the main drivers of energy use and operational costs in activated sludge processes is the amount of oxygen provided to enable biological phosphorus and nitrogen removal. Wastewater treatment facilities are increasingly considering reduced aeration to decrease energy consumption, and whereas successful BNR has been demonstrated in systems with minimal aeration, an adequate understanding of the microbial communities that facilitate nutrient removal under these conditions is still lacking. In this study, we used genome-resolved metagenomics to evaluate the diversity of the “*Candidatus* Accumulibacter” genus in pilot-scale plants operating with minimal aeration. We identified the “*Ca.* Accumulibacter” species enriched under these conditions, including one novel species for which we propose “*Ca.* Accumulibacter jenkinsii” sp. nov. as its designation. Furthermore, the MAGs obtained for 5 additional “*Ca.* Accumulibacter” species further refine the phylogeny of the “*Ca.* Accumulibacter” genus and provide new insight into its diversity within unconventional biological nutrient removal systems.

## INTRODUCTION

Enhanced biological phosphorus removal (EBPR) is a critical process for phosphorus (P) removal from wastewater and helps prevent eutrophication of water bodies receiving the treated effluent (1). EBPR is achieved through polyphosphate accumulating organisms (PAOs), which can intracellularly store polyphosphate. “*Candidatus* Accumulibacter”, commonly referred to as Accumulibacter and affiliated with the *Rhodocyclaceae* family in *Proteobacteria,* has been identified as a PAO commonly found in EBPR systems (2, 3). Accumulibacter possesses the ability to anaerobically generate internal energy and reducing equivalents through the use of intracellularly stored polyphosphate and glycogen. This energy is used to take up volatile fatty acids (VFAs) and convert them to internally stored polyhydroxyalkanoate (PHA). Aerobically, Accumulibacter uses the stored PHA as an energy source for growth and to replenish the intracellularly stored polyphosphate and glycogen (4).

Under these cyclic anaerobic-aerobic conditions, net P removal from the bulk liquid is achieved (5). A key aspect in the design of biological nutrient removal (BNR) treatment plants is the optimization of the anaerobic and aerobic environments that favor Accumulibacter activity (6). However, as BNR treatment plant designs expand towards more energy efficient alternatives such as operation with minimal or no aeration (7-9), the cycles of anaerobic and aerobic conditions often deviate from those initially considered optimal for Accumulibacter. Some members of the Accumulibacter lineage have been shown to thrive on anaerobic and aerobic cycles in which the dissolved oxygen (DO) is severely limited (10). Others thrive under conditions in which the aerobic zone is replaced by an anoxic zone where oxidized nitrogen species serve as electron acceptors (11, 12). The adaptation of Accumulibacter to these different environments illustrates the functional diversity of the lineage (13, 14).

The Accumulibacter lineage has been traditionally subdivided, using the polyphosphate kinase 1 (*ppk1*) gene as a phylogenetic marker, into two types (I and II) and several clades (I-A to I-H and II-A to II-I) (10, 15-18). A recent metagenome-based evaluation of the Accumulibacter lineage proposed the subdivision of the lineage into 18 different “*Ca*. Accumulibacter” species and confirmed the *ppk1*-based classification as a good choice to resolve Accumulibacter phylogeny, as it mirrors genome-based phylogeny (13). Because most metagenome-assembled genomes (MAGs) of Accumulibacter are incomplete and have no assemblage of the rRNA operons, to date it has been difficult to resolve the Accumulibacter phylogeny based on the traditional 16S rRNA gene marker (13).

In this study, we analyzed Accumulibacter MAGs obtained from samples collected from several BNR pilot pants that were operated with reduced aeration (19, 20) and were treating primary effluent under the typical variations in water temperature, flow rates, and loadings that are experienced at full-scale. Half of the recovered Accumulibacter MAGs could be classified as members of five existing “*Ca*. Accumulibacter” species (“*Ca*. A. meliphilus”, “*Ca*. A. delftensis”, “*Ca*. A. propinquus”, “*Ca*. A. contiguus”, and “*Ca*. A. necessarius”). In all cases, the recovered MAGs included fully assembled rRNA operons, allowing us to further refine the 16S rRNA-based phylogenetic analysis of the Accumulibacter lineage and evaluate phylogeny based on the 23S rRNA gene. The other half of the recovered MAGs could not be assigned to existing “*Ca*. Accumulibacter” species. We discovered a cluster of these MAGs that based on *ppk1*, 16S rRNA, 23S rRNA, ribosomal proteins, and metagenome-based phylogenetic analyses formed a coherently separate cluster within Type II Accumulibacter, for which we propose “*Ca*. Accumulibacter jenkinsii” sp. nov. as its designation and a new clade II-J for its placement based on *ppk1* classification. Finally, we used the proposed extended classification and the metagenomic data from the four pilot plants to evaluate the relative abundance of the different members of the Accumulibacter lineage and concluded that regardless of the differences in pilot plant operation, “*Ca*. A. necessarius and “*Ca*. A. propinquus” were the most abundant “*Ca*. Accumulibacter” species in the low-aeration BNR ecosystem.

## RESULTS AND DISCUSSION

### EBPR performance in pilot plants operated with reduced aeration

Four pilot-scale plant experiments were conducted to evaluate BNR performance when aerobic sections of the treatment train are operated with lower DO concentrations than is typically used in conventional BNR operations (Fig. 1). To control aeration, two pilot-scale experiments used ammonia-based aeration control (19) and two experiments used stepwise reduction based on target DO concentrations (20). In one instance (UCTca pilot plant; Fig. 1A), the first 2 out of 3 aerated tanks (UCT3 and UCT4) maintained average continuous DO concentration less than 0.38 mgDO/L by controlling aeration to achieve an ammonium setpoint of 5 mgN/L in the mid-section of the aeration zone (tank UCT4). The last aerated tank (UCT5) was operated to maintain a DO concentration of 1.0 mgDO/L (19). In a second treatment train (AOia; Fig. 1B), an intermittent aeration scheme was implemented in the aerated zone, with aeration in the first 3 tanks (tanks AO2, AO3, and AO4) controlled by allowing cyclic accumulation and consumption of ammonium. Aeration to the first 3 aerated tanks (AO2, AO3, AO4) was turned on when the ammonium concentration in tank AO4 reached 5 mgN/L and turned off when the ammonium in the same tank reached 2 mgN/L. During the aerated periods, DO was not allowed to accumulate above 0.7 mgDO/L. The last aerated tank of this treatment train (AO5) was a high-DO polishing step in which the DO was constantly maintained at 2.0 mgDO/L. This ammonia-controlled aeration scheme resulted in aeration being off 42% of the time in the aerated tanks (19). In the third and fourth pilot-scale experiments (AO-G, Fig. 1C, and AO-FF, Fig. 1D) aeration was controlled by setting different DO setpoints in the first three aerated tanks and using the last aerated tank as a high-DO polishing step. The first aerated tank in each treatment train was started with a setpoint of 0.4 mgDO/L and was gradually reduced to 0.2 mgDO/L, the second aerated tank started with a setpoint of 0.8 mgDO/L and had stepwise reductions to 0.3 mgDO/L, the third aerated tank was initiated with a setpoint of 3.5 mgDO/L and gradually reduced to 0.5 mgDO/L. The last aerated tank was maintained at 3.5 mgDO/L (20).

**Figure 1.**
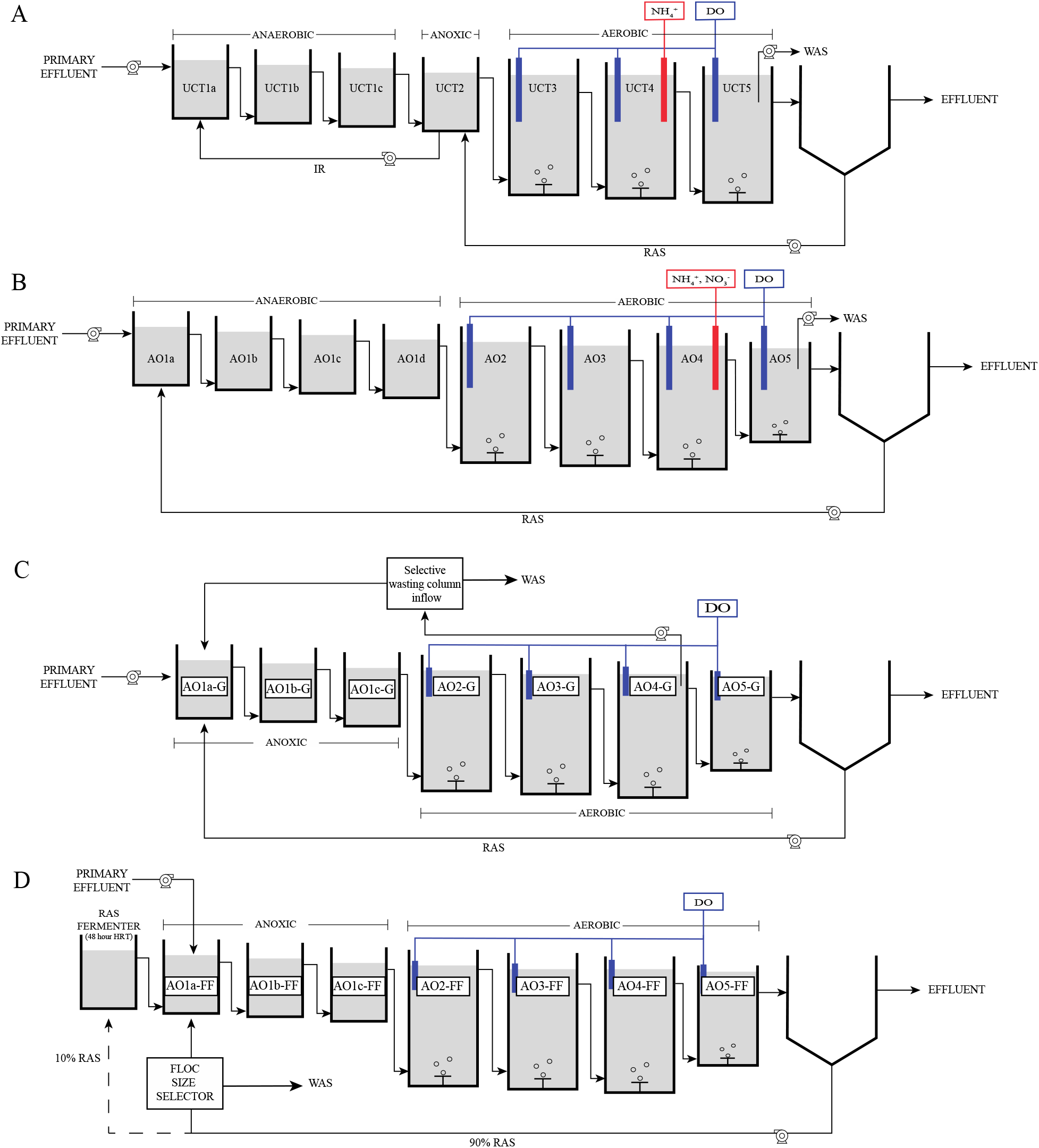
Configurations of the four pilot-scale biological nutrient removal (BNR) treatment plants operated with reduced aeration. (A) UCTca, (B) AOia, (C) AO-G and (D) AO-FF. Dissolved oxygen (DO) sensors in the aerated tanks of all four pilot plants, and ammonium sensors in the UCTca and AOia pilot plants were used to control the amount of air delivered to each reactor. Abbreviations: RAS, return activated sludge; WAS, waste activated sludge; IR, internal recycle; HRT, hydraulic retention time

All treatment trains demonstrated efficient P removal regardless of the aeration mode used (Table 1). Influent filtered total P (TP) to the first set of pilot-scale experiments (UCTca and AOia) and the second set of experiments was 4.3 ± 1.9 and 5.5 ± 1.4 mgP/L, respectively. Average effluent filtered TP concentration in the treatment trains ranged from 0.3 mgP/L to 0.4 mgP/L.

**Table 1.** Average nutrient removal in the four pilot-scale systems.

| Pilot Plant | Average Influent<sup>a,b</sup> |  | Average Effluent<sup>a</sup> |  |  |  |
| --- | --- | --- | --- | --- | --- | --- |
|  | TP<br>(mgP/L) | TKN<br>(mgN/L) | TP<br>(mgP/L) | TKN<br>(mgN/L) | NO<sub>3</sub><sup>-</sup><br>(mgN/L) | NO<sub>2</sub><sup>-</sup><br>(mgN/L) |
| UCTca | 4.3 ± 1.9 | 38.6 ± 8.03 | 0.4 ± 0.6 | 2.1 ± 1.0 | 13.0 ± 3.9 | 0.3 ± 0.3 |
| AOia |  |  | 0.3 ± 0.4 | 1.8 ± 1.1 | 11.2 ± 3.2 | 0.2 ± 0.2 |
| AO-G | 5.5 ± 1.4 | 46.2 ± 6.3 | 0.4 ± 0.7 | 2.4 ± 3.0 | 15.8 ± 4.1 | 0.8 ± 1.9 |
| AO-FF |  |  | 0.3 ± 0.6 | 4.4 ± 6.9 | 14.5 ± 6.1 | 0.6 ± 1.0 |

<sup>a</sup>Averages of influent and effluent filtered samples. Abbreviations: TP, total phosphorus; TKN, total Kjeldahl nitrogen; NO<sub>3</sub><sup>-</sup>, nitrate; NO<sub>2</sub><sup>-</sup>, nitrite.
<sup>b</sup>Pilot plants UCTca and AOia were operated in parallel and received the same influent. Likewise, Pilot plants AO-G and AO-FF were operated in parallel and received the same effluent.

In addition to P removal, BNR processes are operated to also achieve nitrogen removal (21). Average filtered influent total Kjheldal nitrogen (TKN) was 38.6 ± 8.0 and 46.2 ± 6.3 mgTKN/L to the first and second set of pilot plant experiments, respectively, and average effluent filtered TKN ranged between 1.8 and 4.4 mgTKN/L in the four pilot plants (Table 1), demonstrating efficient nitrification under the reduced aeration conditions. No substantial nitrite (NO_2_^-^) accumulation was observed in the effluent of any of the treatment trains (averages were less than 1.0 mgNO_2_^-^-N/L). The pilot-scale systems were not configured to maximize total nitrogen removal (Fig. 1) since they mimicked full-scale operation at the Nine Springs wastewater treatment plant (Madison, WI), which is a process that does not include internal nitrate recycle (22). Under the reduced aeration conditions, effluent nitrate (NO_3_^-^) ranged from 11.2 to 15.8 mgNO_3_^-^-N/L (Table 1).

### Assembly and Classification of “*Ca*. Accumulibacter” MAGs

Having determined that efficient EBPR was established in the four pilot plants operated with reduced aeration, we proceeded to analyze the microbial communities at different times during pilot plant operations. In total, we obtained individual metagenomes from 15 6-10kb PacBio, 7 3kb PacBio, and 2 Illumina libraries (Table S1). Among the MAGs assembled from these metagenomes were 16 MAGs taxonomically classified, according to the GTDB-Tk Lineage classification, as belonging to the Accumulibacter lineage. We named these MAGs using strain names UW14 to UW29 (Table 2), adding to the UW# strain designations in the collection of “*Ca*. Accumulibacter” MAGs that have been assembled from activated sludge originally collected from the Madison Metropolitan Sewerage District wastewater treatment plant (23-27).

**Table 2.**
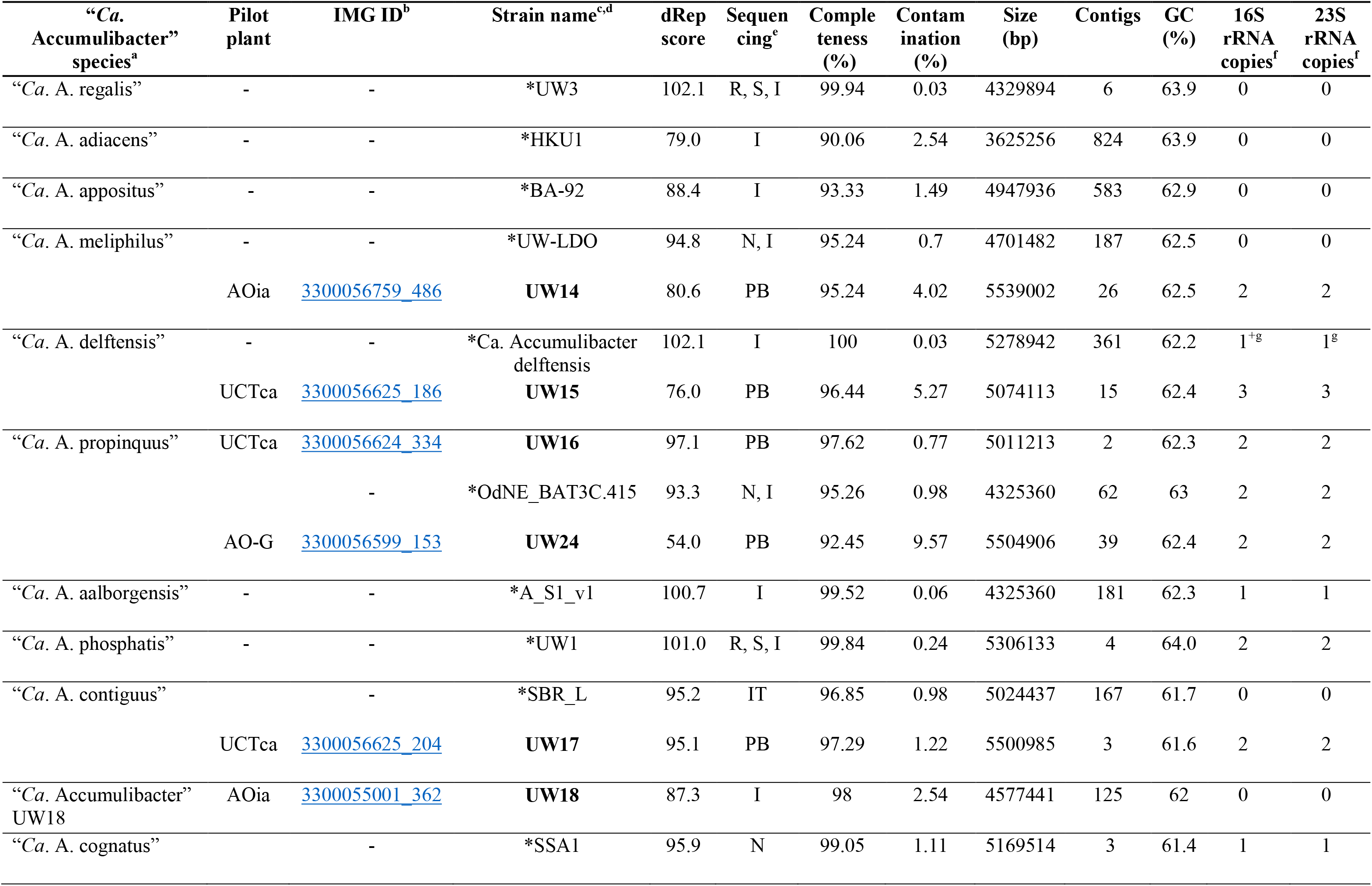
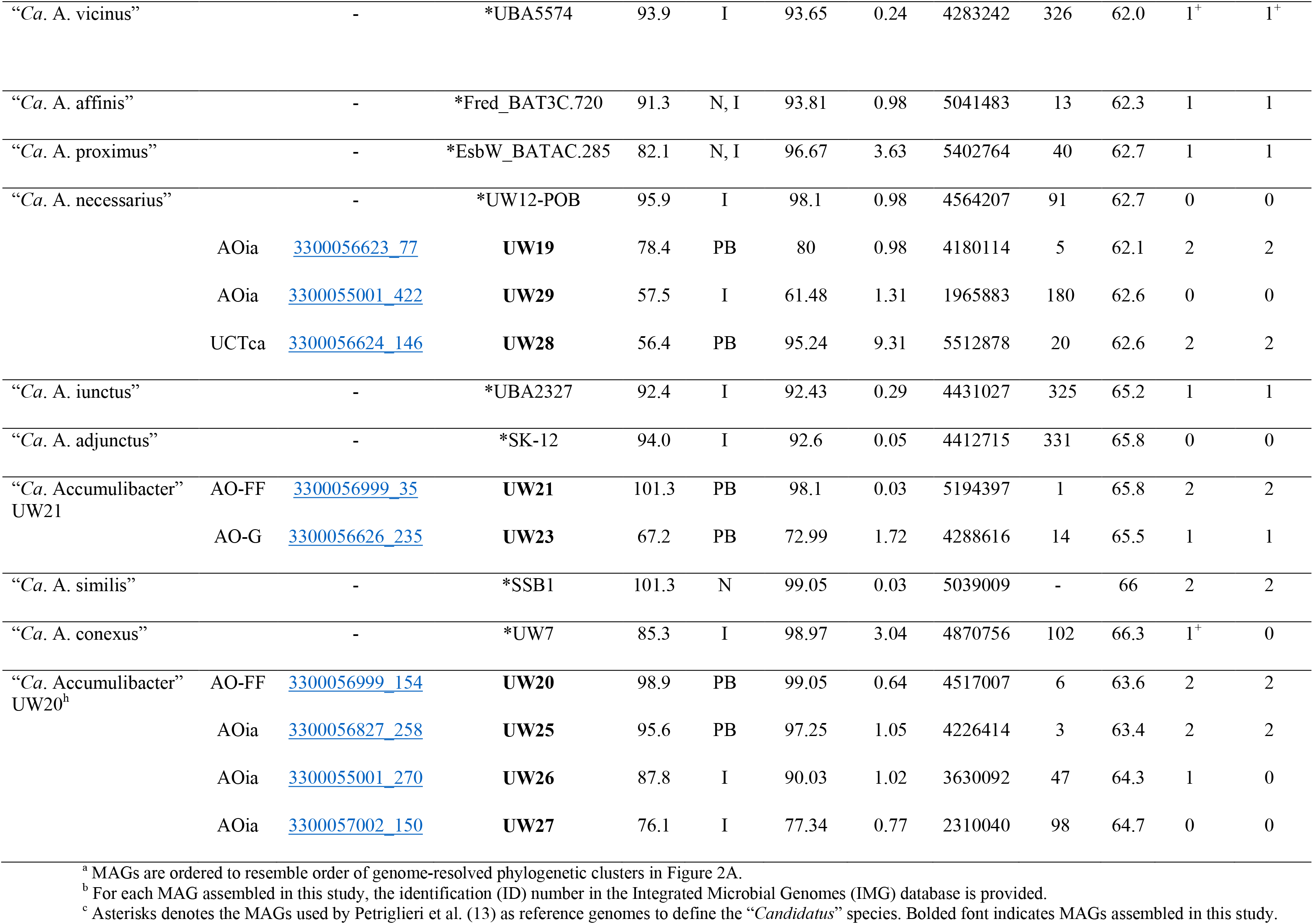
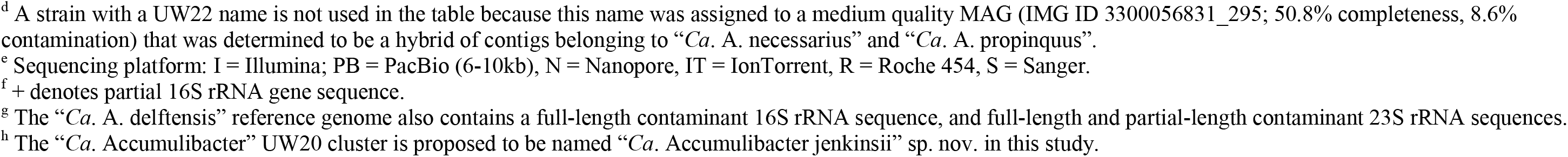
Statistics of MAGs and reference genomes for each “*Ca*. Accumulibacter” species.

To classify these Accumulibacter MAGs according to the established “*Ca*. Accumulibacter” taxonomy, we used them along with the reference genomes for each “*Ca*. Accumulibacter” species, as defined by Petriglieri et al. (13), to obtain genome clusters belonging to each species (Fig. S1). This analysis revealed that 8 out of the 16 assembled MAGs could be classified as belonging to 5 different “*Ca*. Accumulibacter” species since they had ANI values greater than 95% in pairwise comparisons with the reference genome for the species (Fig. S2). That is, UW14 belonged to “*Ca*. A. meliphilus,” UW15 to “*Ca*. A. delftensis,” UW16 and UW24 to “*Ca*. A. propinquus,” UW17 to “*Ca*. A. contiguus,” and UW19, UW28, and UW29 to “*Ca*. A. necessarius” (Table 2).

A group of four genomes (UW20, UW25, UW26, and UW27) formed a separate cluster based on dRep’s primary Mash clustering (Fig. S1), with UW20 having the highest quality according to the dRep score, and thus, we used UW20 as the initial name for this cluster (Table 2). Two of the genomes in this cluster, UW20 and UW25 were obtained from PacBio libraries and are of high quality (HQ, greater than 90% completeness, less than 5% contamination) according to MiMAG standards (28). In addition, they have two fully assembled copies of the rRNA operon, which facilitates additional analysis of this novel cluster and the proposal for a new species epithet (see below).

Two additional genomes (UW21 and UW23) also formed a separate cluster (Fig. S1), with UW21 being an HQ MAG with fully assembled rRNA operons. This cluster was closely associated with “*Ca*. A. adjunctus”, with ANI values just below the threshold of 95% used for species designations (ANI 94.7 and 94.6% for UW21 and UW23, respectively). The incomplete reference genome for “*Ca*. A. adjunctus” does not include 16S or 23S rRNA sequences, and therefore, a complete assessment of whether the genomes in this cluster represent strains belonging to “*Ca*. A. adjunctus” cannot be made. Thus, we designate this cluster as “Ca. Accumulibacter” UW21 (Table 2).

An additional MAG assembled from an Illumina library, UW18, also formed a separate cluster at the 95% ANI threshold (Fig. S1). The closest reference genomes were those of “*Ca*. A. vicinus (ANI 91.8%) and “*Ca*. A. cognatus” (ANI 91.5%). This is also an HQ genome, but the assembly did not include the rRNA operon (Table 2). We designate this cluster as “Ca. Accumulibacter” UW18.

Finally, there was an additional medium-quality genome (UW22, 50.9% completeness and 8.6% contamination) assembled from a PacBio library that also clustered outside of the established “*Ca*. Accumulibacter” species (Fig. S1). This MAG contained 4 assembled copies of the rRNA operon. Upon further inspection of the contigs in this MAG, and of the 16S rRNA sequences (see below), we determined that this MAG was a hybrid, containing a combination of contigs from “*Ca*. A. propinquus” and “*Ca*. A. necessarius.” Therefore, this MAG was removed from further analysis, except for the rRNA sequences, which provided useful information in the rRNA-based phylogenetic analysis.

### Accumulibacter phylogeny based on *ppk1*, genome-scale, and ribosomal proteins

The diversity of the Accumulibacter lineage has typically been described using a *ppk1*-based phylogeny, which subdivides the lineage into two Types and several phylogenetically coherent clades. In the most comprehensive classification to date (13, 25), eight clades have been defined for Type I (I-A to I-H) and nine clades for Type II (II-A to II-I). However, representative genomes are only available for three clades within Type I (I-A, I-B, and I-C) and six clades within Type II (II-A, II-B, II-C, II-D, II-F, and II-G), and thus, the defined “*Ca*. Accumulibacter” species belong to these 9 clades. For the “*Ca*. Accumulibacter” species there is also agreement in the phylogenies obtained from the analysis of *ppk1* sequences and genome-based phylogenies usually created from the concatenated alignment of single copy marker genes (13).

We compared the *ppk1*-based phylogenies to the genome-base phylogenies for the “*Ca*. Accumulibacter” reference genomes and the UW14 to UW29 Accumulibacter MAGs (Fig. 2) and found the same agreement in phylogenies as reported by Petriglieri et al. (13). Moreover, the phylogenetic trees place the UW20 cluster in separate branches, and based on the *ppk1*-based tree, the *ppk1* sequences in the UW20 cluster belong to Type II, but do not cluster with any of the clades that include the recognized “*Ca*. Accumulibacter” species (Fig. 2A). Furthermore, extending the *ppk1* analysis to include sequences that define the clades not represented by any of the “*Ca*. Accumulibacter” species show that the *ppk1* gene sequences of the UW20 cluster do not cluster with any of the other defined clades (Fig. S3), indicating that the UW20 cluster defines a novel clade within the *ppk1*-based Type II lineage. Thus, we propose a definition of a new Type II clade, clade II-J, to identify the “Ca. Accumulibacter” UW20 cluster.

**Figure 2.**
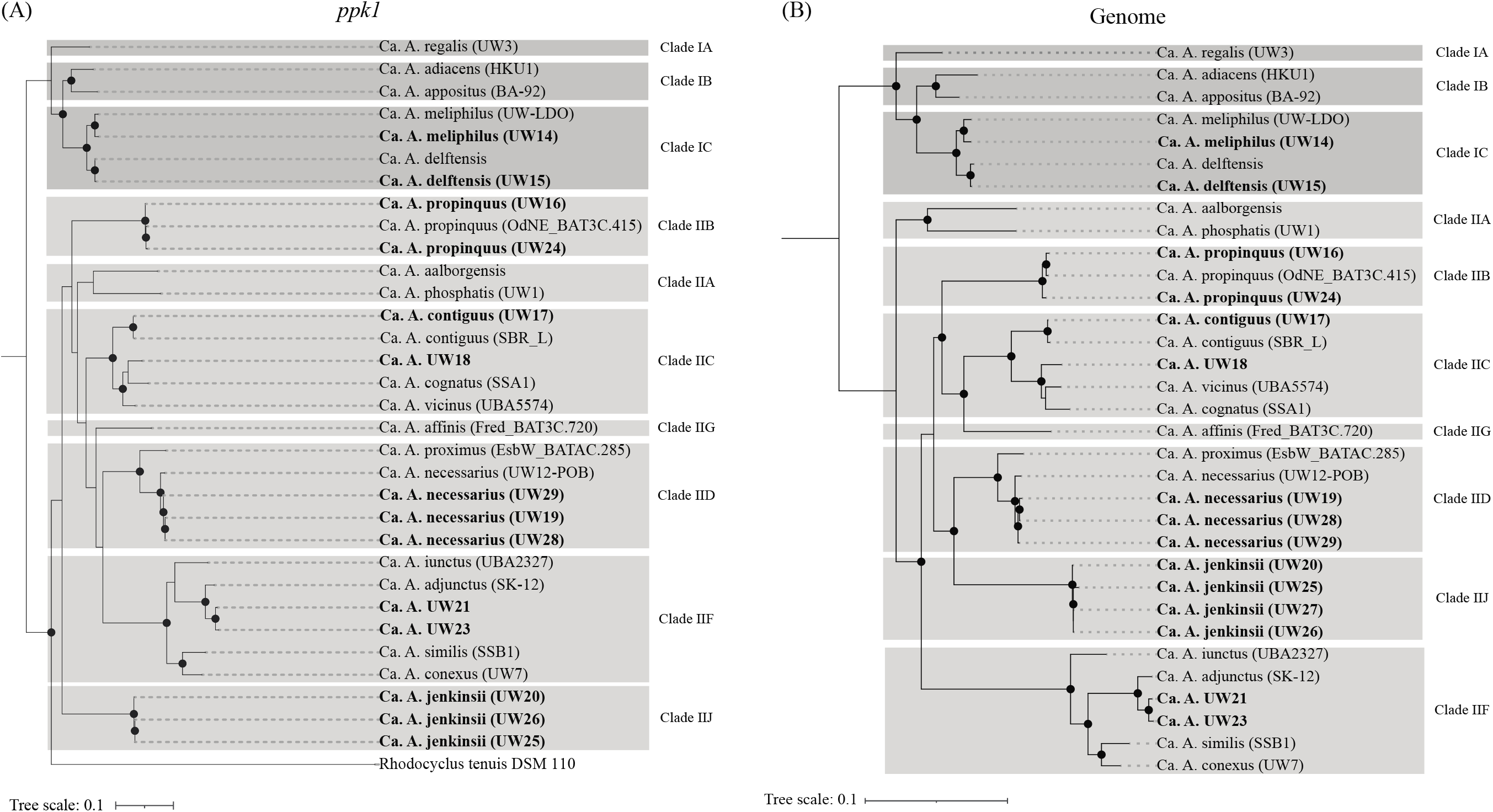
Phylogenetic analyses of reference “Ca. Accumulibacter” species and Accumulibacter MAGs assembled in this study (bolded tips) based on (A) complete nucleotide sequence of the *ppk1* gene, and (B) the alignment of 120 single-copy conserved marker genes identified with GTDB-Tk. Each tree was constructed in RAxML-NG using 100 bootstraps. Nodes with bootstraps greater than 70 are denoted with black circles. Grey boxes indicate clade designations based on *ppk1* phylogeny. The *ppk1* gene was not present in the incomplete assembly of UW27, and UW22 was excluded from both trees due to the MAG being deemed to be a hybrid of different species.

An additional recently proposed genome-based phylogenetic analysis to classify members of the Accumulibacter lineage uses the alignment of 16 concatenated ribosomal proteins (29). We performed this analysis with the reference genomes for the “*Ca*. Accumulibacter” species plus the UW14 to UW29 Accumulibacter MAGs (excluding UW22) for which a complete set of these ribosomal proteins was included in the assembled metagenomes (Fig. 3). The phylogeny from this analysis also resembles the *ppk1*-based classification of the Accumulibacter lineage and separates “*Ca*. Accumulibacter” species into two Types and multiple clades within each Type. Furthermore, it also supports the placing of the UW20 cluster in a separate clade within Type II (i.e., Type II-J).

**Figure 3.**
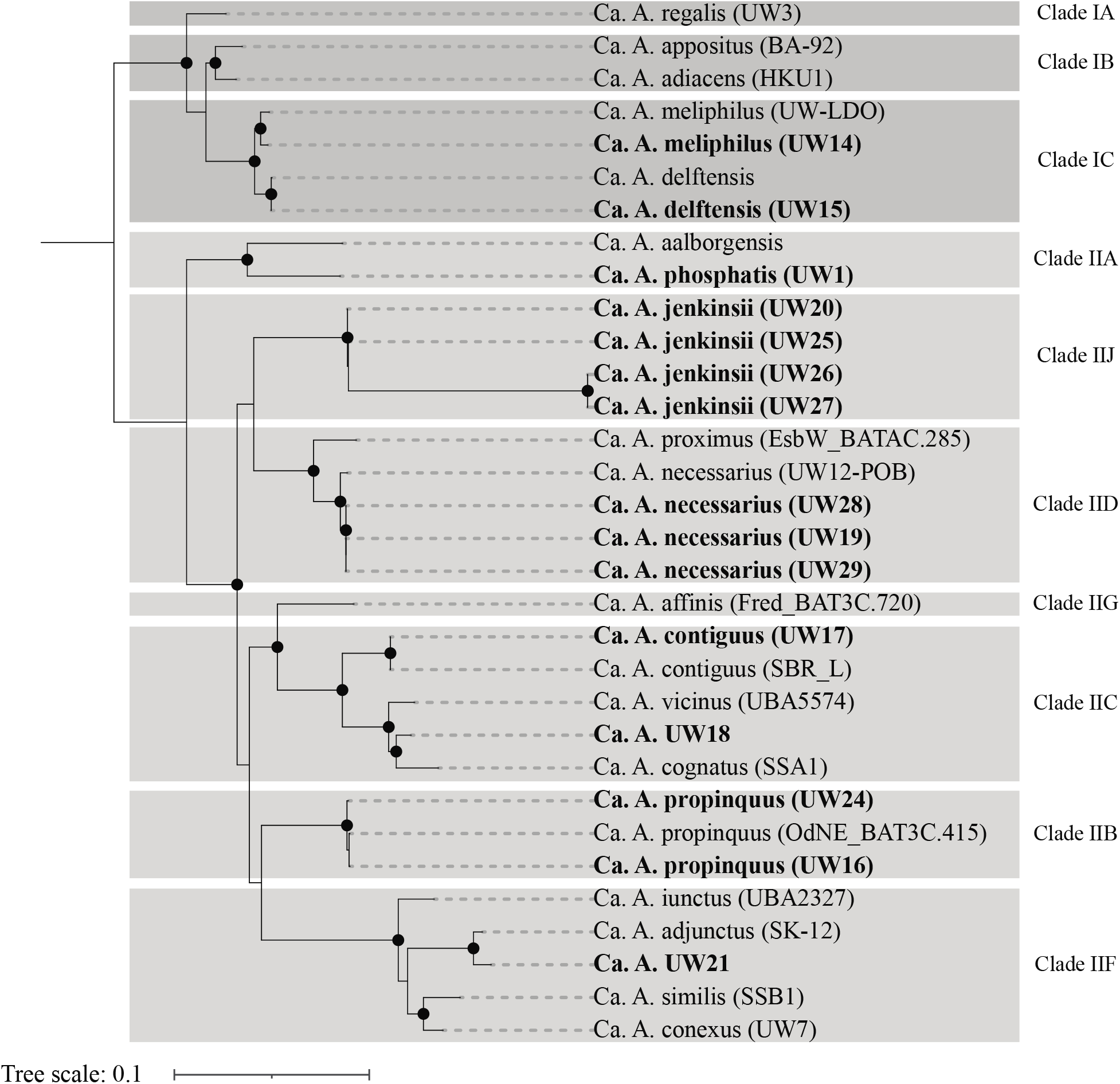
Concatenated ribosomal protein tree of the UW14 to UW29 Accumulibacter MAGs, along with reference genomes for the “*Ca*. Accumulibacter” species, using a set of 16 ribosomal marker proteins. The tree was constructed in RAxML with 100 bootstraps. Nodes with bootstraps greater than 70 are denoted with black circles. Bolded tips represent Accumulibacter MAGs assembled in this study. The incomplete assembly of UW23 did not include a complete set of marker proteins and is not included. UW22 is not included because this MAG was deemed to be a species hybrid.

### Reassessment of “*Ca*. Accumulibacter” phylogeny based on rRNA gene sequences

The 16S rRNA-based phylogeny of “*Ca*. Accumulibacter” is generally not considered to provide an adequate representation of the diversity of this genus because it cannot resolve the genome-inferred or *ppk1*-inferred differences (13, 30, 31). This may be in part because many of the assembled metagenomes of “*Ca*. Accumulibacter” do not include complete copies of the rRNA operons. Indeed, after the recent proposal of new “*Ca*. Accumulibacter” species by Petriglieri et al. (13), the reference genomes for only six “*Candidatus*” species contained full-length 16S rRNA sequences (“*Ca*. A. phosphatis”, “*Ca*. A. propinquus”, “*Ca*. A. affinis”, “*Ca*. A. proximus”, “*Ca*. A. iunctus”, and “*Ca*. A. similis”). Furthermore, out of the “*Ca*. Accumulibacter” species that had been proposed earlier (23, 32, 33), “*Ca*. A. phosphatis” and “*Ca*. A. aalborgensis” possessed full 16S rRNA gene sequences, but “*Ca*. A. delftensis” was assembled with only a partial 16S rRNA gene sequence that matches the Accumulibacter lineage and a contaminant 16S rRNA sequence that does not belong to the Accumulibacter lineage (and excluded from this analysis). A recent analysis of the reference genomes available at NCBI for other “*Ca*. Accumulibacter” species shows full-length 16S rRNA gene sequence for “*Ca*. A. cognatus” and partial sequences for “*Ca*. A. vicinus” and “*Ca*. A. conexus” (Table 2).

The MAGs in this study provide additional full-length 16S rRNA gene sequences for four “*Ca*. Accumulibacter” species for which their reference genomes lack full-length 16S rRNA sequences: “*Ca*. A. meliphilus”, “*Ca*. A. delftensis”, “*Ca*. A. contiguus”, and “*Ca*. A. necessarius”. In addition, the “*Ca*. Accumulibacter” clusters UW20 and UW21 possess complete rRNA operons assembled.

Thus, with the additional information provided by the UW14 to UW29 metagenomes, 13 out of the 18 “*Ca*. Accumulibacter” species possess full-length 16S rRNA sequences, and two have partial 16S rRNA sequences. This allows for a more comprehensive analysis of the “*Ca*. Accumulibacter” phylogeny based on the 16S rRNA gene (Fig. 4). For this analysis, we aligned the 16S rRNA sequences available for reference genomes of “*Ca*. Accumulibacter” species and the 16S rRNA sequences extracted from the “*Ca*. Accumulibacter” MAGs assembled in this study. The information from the reference genomes and “*Ca*. Accumulibacter” MAGs were complemented with Accumulibacter-related 16S rRNA sequences that were extracted from fully assembled rRNA operons in a metagenome assembly that did not result in the binning of an Accumulibacter-related MAG (Ga0558717; Table S1), and for phylogenetic context, the MiDAS4 full-length sequences from amplicon sequence variants (ASV) assigned to the Accumulibacter lineage included in the analysis of Petriglieri et al. (13).

**Figure 4.**
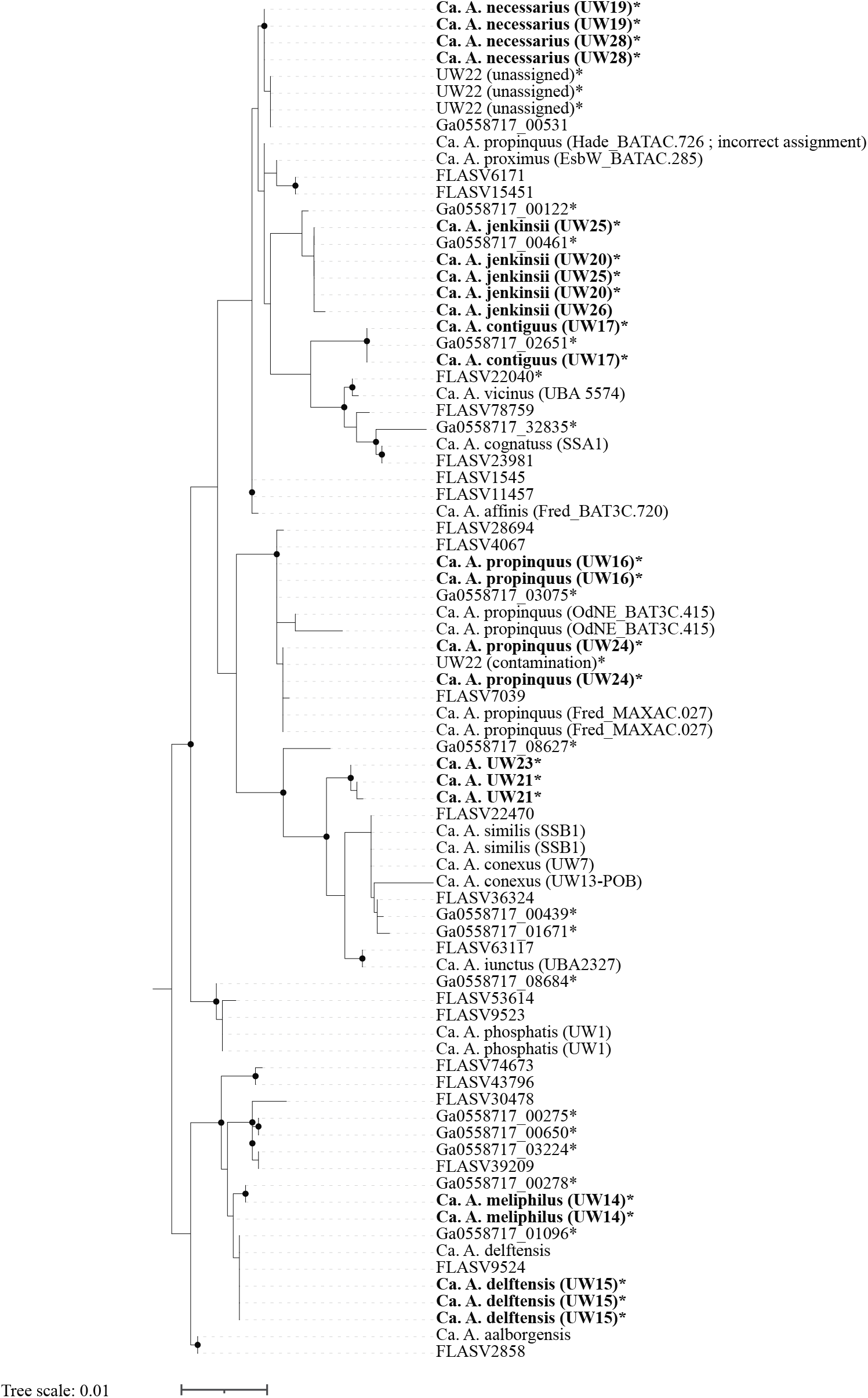
Phylogenetic tree of aligned 16S rRNA gene sequences retrieved from assembled metagenomes from this study (UW14 to UW29; bolded tips), reference genomes for the “Ca. Accumulibacter” species, the MiDAS database (FLASV sequences), and from complete rRNA operons obtained from one of the assemblies (Ga0558717 sequences). The tree was constructed in RAxML-NG using 1,000 bootstraps. Starred tips indicate sequences from this study found in complete operons and for which the corresponding 23S rRNA sequence was included in the phylogenetic tree in Figure 5.

Because the long PacBio reads facilitate assemblies of complete rRNA operons, we also constructed a 23S rRNA tree (Fig. 5) that included sequences from the reference genomes, the new “*Ca*. Accumulibacter” MAGs, and other fully assembled operons found in the assembly and for which the corresponding 16S rRNA sequence was included in the 16S rRNA tree. The reference genomes for “*Ca*. A. aalborgensis”, “*Ca*. A. propinquus”, “*Ca*. A. cognatus”, “*Ca*. A. affinis”, “*Ca*. A. proximus”, “*Ca*. A. iunctus”, and “*Ca*. A. similis” possess full-length 23S rRNA gene sequences. The reference genome for “*Ca*. A. delftensis” includes three full-length 23S rRNA gene sequences, two of which are contaminants not related to the Accumulibacter lineage and excluded from the analysis. The MAGs assembled in this study add full-length 23S rRNA gene sequences for “*Ca*. A. meliphilus”, “*Ca*. A. delftensis”, “*Ca*. A. contiguus”, “*Ca*. A. necessarius”, and the UW20 and UW21 clusters.

**Figure 5.**
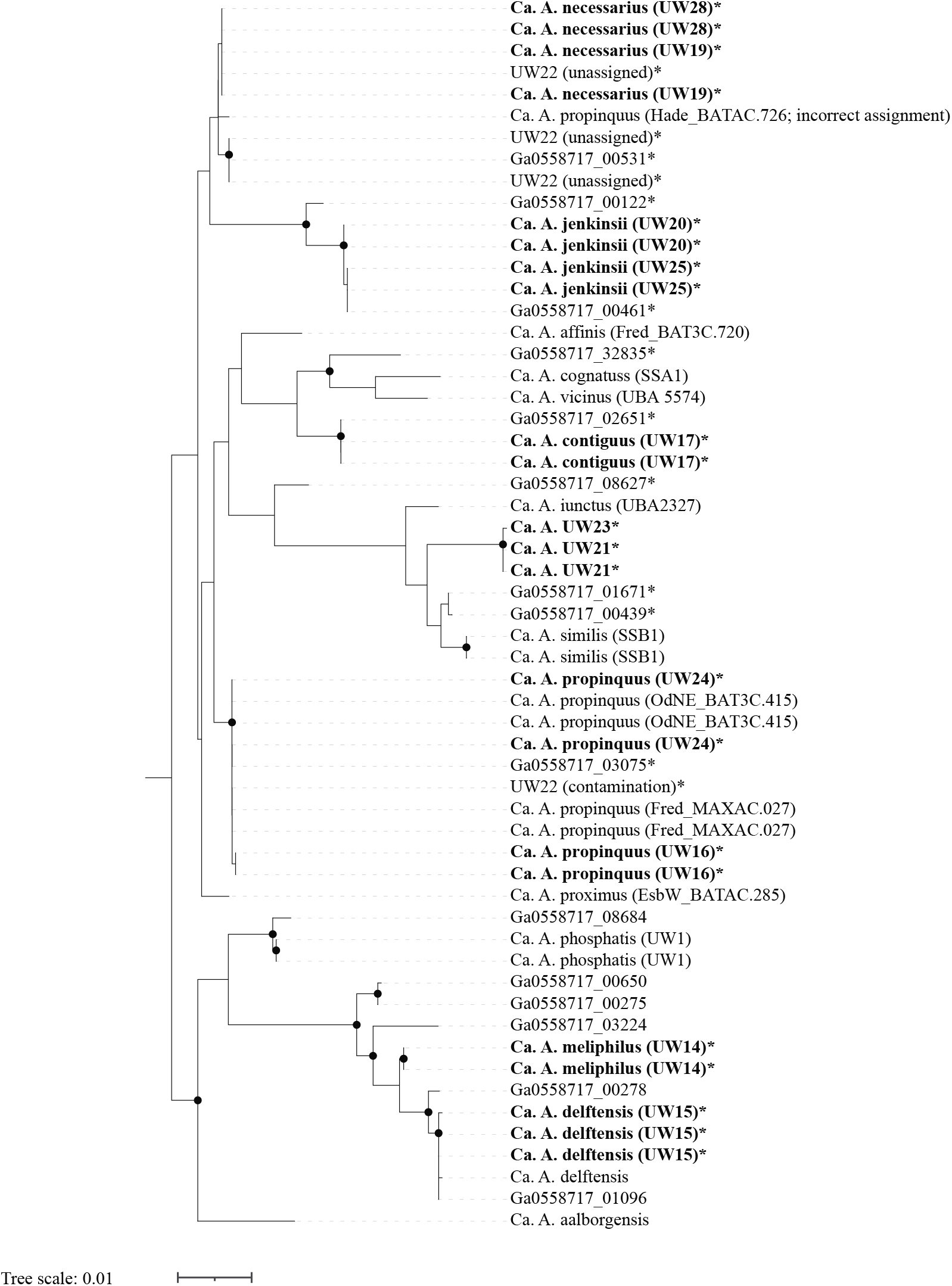
Phylogenetic tree of aligned 23S rRNA gene sequences retrieved from Accumulibacter MAGs and assembled metagenomes from this study, the reference genomes of the “*Ca.* Accumulibacter” species, and from complete rRNA operons obtained from one of the assemblies (Ga0558717 sequences). The tree was constructed in RAxML-NG using 1,000 bootstraps. Bolded tips represent 23S rRNA sequences recovered from MAGs assembled in this study, and starred tips indicate sequences from complete rRNA operons for which the corresponding 16S rRNA sequence was included in the phylogenetic tree in Figure 4.

As anticipated from prior evaluations (13), the 16S rRNA analysis did not sufficiently resolve all “*Ca*. Accumulibacter” species (Fig. 4). The 23S rRNA-based tree (Fig. 5) showed general agreement with the 16S rRNA-based tree. In the two trees, some “*Ca*. Accumulibacter” species form their own separate clusters and others were not sufficiently resolved. The resolved clusters generally reflect those that emerge from the *ppk1*-based phylogeny (Fig. 2).

New in this analysis are the 16S and 23S rRNA sequences for “*Ca*. A. meliphilus” (UW14) and “*Ca*. A. delftensis” (UW15). The 16S rRNA sequences from these MAGs clustered together in both trees along with the partial 16S rRNA and one complete 23S rRNA sequence found in the reference genome of “*Ca*. A. delftensis”, respectively. In addition, five pairs of 16S and 23S sequences found in complete operons in other contigs in the assembly also belonged to these clusters, confirming the agreement between the clustering in the 16S and 23S rRNA-based trees. These two “*Ca*. Accumulibacter” species are currently the only ones defined as belonging to the *ppk1*-based clade I-C, so the clustering of sequences of these two species together in the rRNA-based trees indicates agreement with *ppk1*-based phylogeny, although they cannot be properly differentiated from each other based on rRNA sequence analyses.

Also new in this analysis are the 16S and 23S rRNA sequences for “*Ca*. A. contiguus” (strain UW17). In both trees, these sequences formed their own separate sub-cluster within a cluster that also included the sequences from the reference genomes of “*Ca*. A. vicinus” and “*Ca*. A. cognatus”, species for which we did not recover MAGs in this study. One additional pair of 16S and 23S sequences obtained from a complete rRNA operon (Ga0558717_02651 in Figs. 4 and 5) had identical sequences to the two operons found in UW17, further supporting the correct assembly of these operons in UW17. Thus, we conclude that the new rRNA sequences for “*Ca*. A. contiguus” shows that the rRNA-based clustering agrees with the *ppk1*-based phylogeny since “*Ca*. A. contiguus”, “*Ca*. A. vicinus” and “*Ca*. A. cognatus” are the only three species currently defined as belonging to clade II-C.

This study also produced new 16S and 23S rRNA sequences for “*Ca*. A. necessarius”, a species for which there were no available rRNA sequences in the reference genome (13). Two complete rRNA operons were assembled for UW19 and UW28, MAGs that were classified as “*Ca*. A. necessarius” by ANI (Table 2). The rRNA sequences consistently formed their own separate cluster along with the sequences from three operons assembled in UW22 and sequences from another operon found in an assembly contig that was not binned (Ga0558717_00531). A fourth rRNA operon that was assembled into UW22 did not cluster with the rest of the UW22 sequences (referred to as “UW22 (contamination)” in Figs. 4 and 5), instead clustering with sequences of “*Ca*. A. propinquus”. The unusual observation of four rRNA operons in a single “*Ca*. Accumulibacter” MAG, and the fact that one of the operons had sequences that did not cluster with the other three, are strong evidence to indicate that MAG UW22 is a chimeric assembly (we did not include UW22 in Table 2). It is surprising to see that the 16S and 23S sequences for strain MAXAC.726, a MAG reclassified as “*Ca*. A. propinquus” by Petriglieri et al. (13), did cluster near the “*Ca*. A. necessarius” sequences instead of the “*Ca*. A. propinquus” sequences. This possibly indicates contamination in the assembly of the MAXAC.726 MAG.

The MAGs UW16 and UW24 were classified as belonging to “*Ca*. A. propinquus” by ANI comparisons (Table 2). Each of these MAGs has two fully assembled rRNA operons, and consistent with the ANI classification, the 16S and 23S rRNA sequences clustered with the sequences from the reference genome for this species (strain BAT3C.415) and with the sequences of another genome reclassified by Petriglieri et al. (13) as also belonging to this species (strain MAXAC.027). “*Ca*. A. propinquus” is the only “*Ca*. Accumulibacter” species in clade II-B, so the 16S and 23S sequences forming their own separate cluster is consistent with the *ppk1*-based classification.

The 16S rRNA sequences from UW21 and UW23 clustered with sequences from the reference genomes of “*Ca*. A. iunctus”, “*Ca*. A. similis”, and “*Ca*. A. conexus”, three out of the four “*Ca*. Accumulibacter” species defined as belonging to clade II-F. “*Ca*. A. adjunctus” is the fourth species in this clade, but its reference genome does not have 16S rRNA sequences assembled. In the 23S rRNA tree, the sequences of UW21 and UW23 clustered together with the 23S rRNA sequences of “*Ca*. A. iunctus”, “*Ca*. A. similis”, the only two reference genomes in this clade that had 23S rRNA sequences assembled. Thus, the rRNA analyses would place the UW21 cluster (UW21 and UW23 MAGs; Table 2) as members of the *ppk1*-based clade II-F, in agreement with the *ppk1* and genome-based analyses (Fig. 2). In the phylogenetic tree based on ribosomal proteins (Fig. 3), the UW21 MAG also clustered in clade II-F, whereas the UW23 MAG was not included in this analysis due to having an incomplete set of ribosomal proteins.

Sequences from the UW20 cluster MAGs (UW20, UW25, and UW26) formed independent clusters in both the 16S and the 23S rRNA trees, in agreement with the other phylogenetic analyses (Figs. 2 and 3) that suggest that the MAGs in this cluster represent a yet undescribed “*Ca*. Accumulibacter” species.

In a final analysis of the 16S rRNA sequences, we performed an *in silico* analysis of the DNA probes that are used in fluorescence *in situ* hybridization (FISH) to visualize Accumulibacter cells (Table 3). Probe Acc470, designed to target “*Ca*. A. delftensis” (13) is a perfect match to UW15 (“*Ca*. A. delftensis”) and to UW14 (“*Ca*. A. meliphilus”), expanding the number of species that this probe targets, and showing agreement with the difficulty to differentiate these two species by their 16S rRNA sequences (Fig. 4). Consistent with the placement of UW16 and UW24 as members of the “*Ca*. A. propinquus” species, the probe designed to visualize “*Ca*. A. propinquus” by Petriglieri et al. (13) is also a perfect match to these two MAGs. The new 16S rRNA sequences for “*Ca*. A. contiguus” showed mismatches to all the species-specific probes, but with only one mismatch to the “*Ca*. A. proximus” Acc469 probe, the application of Acc469 may require an additional competitor probe. Likewise, the new 16S rRNA sequences for “*Ca*. A. necessarius” shows only one mismatch with the Acc469 probe and with the Acc635 probe targeting “*Ca*. A. regalis”. The UW21 cluster shows perfect matches to probes Acc471 and Acc471_2, complicating the application of these two probes that target the same site in the 16S rRNA sequence but aim at targeting two different groups of “Ca. Accumulibacter” species (Table 3). Finally, the 16S rRNA sequences of cluster UW20 show two or three mismatches to all the species-specific probes.

**Table 3.** Mismatch analysis of FISH probes to new 16S rRNA sequences obtained in this study for “*Ca*. Accumulibacter” species.

| “*Ca. Accumulibacter*”<br>species | Strain<br>name | Type<sup>a</sup> | PAO651<sup>b</sup><br>PAO<br>cluster | Acc-I-444<sup>c</sup><br>Type I | Acc-II-444<sup>c</sup><br>Type II | Acc469<sup>d</sup><br>“*Ca. A.*<br>*proximus*” | Acc470<sup>d</sup><br>“*Ca. A.*<br>*aalborgensis*”<br>and “*Ca. A.*<br>*delftensis*” | Acc471<sup>d</sup><br>“*Ca. A.*<br>*affinis*” and<br>“*Ca. A.*<br>*proximus*” | Acc471_2<sup>d</sup><br>“*Ca. A.*<br>*iunctus*”<br>and “*Ca. A.*<br>*similis*” | Acc635<sup>d</sup><br>“*Ca. A.*<br>*regalis*” | Acc1011<sup>d</sup><br>“*Ca. A.*<br>*propinquus*” |
| --- | --- | --- | --- | --- | --- | --- | --- | --- | --- | --- | --- |
| “*Ca. A. meliphilus*” | UW14 | I | PM<sup>b</sup> | PM | 2 | 4 | PM | 2 | 2 | 2 | 1 |
| “*Ca. A. delftensis*” | UW15 | I | PM | PM | 2 | 4 | PM | 2 | 2 | 2 | 1 |
| “*Ca. A. propinquus*” | UW16 | II | PM | 3 | 3 | 4 | 3 | 3 | 3 | 1 | PM |
|  | UW24 | II | PM | 3 | 3 | 4 | 3 | 3 | 3 | 1 | PM |
| “*Ca. A. contiguus*” | UW17 | II | PM | 2 | PM | 1 | 3 | 4 | 4 | 2 | 3 |
| “*Ca. A. necessarius*” | UW19 | II | PM | 1 | 1 | 1 | 3 | 4 | 4 | 1 | 2 |
|  | UW28 | II | PM | 1 | 1 | 1 | 3 | 4 | 4 | 1 | 2 |
| “*Ca. Accumulibacter*”<br>UW20<sup>e</sup> | UW20 | II | PM | 2 | PM | 2 | 2 | 3 | 3 | 2 | 2 |
|  | UW25 | II | PM | 2 | PM | 2 | 2 | 3 | 3 | 2 | 2 |
| “*Ca. Accumulibacter*”<br>UW21 | UW21 | II | 1 | 2 | PM | 3 | 2 | PM | PM | 2 | 1 |
|  | UW23 | II | 1 | 2 | PM | 3 | 2 | PM | PM | 2 | 1 |

<sup>a</sup> Phylogenetic assignment of Type I and Type II based on *ppk1* gene sequences.
<sup>b</sup> Probe originally reported by Crocetti et al. (57)
<sup>c</sup> Probe originally reported by Flowers et al. (58)
<sup>d</sup> Probe originally reported by Petriglieri et al. (13)
<sup>e</sup> The “*Ca. Accumulibacter*” UW20 cluster is proposed to be named “*Ca. Accumulibacter jenkinsii*” sp. nov. in this study.

### Phylogenetic classification of the UW20 cluster

All phylogenetic analyses performed, based on 16S rRNA, 23S rRNA, *ppk1*, ribosomal proteins, and genome-based sequence alignments consistently support the UW20 cluster as representing a new species within the Accumulibacter lineage. Four different MAGs were assembled in this cluster, from independent metagenome libraries (Table 2) that originated from two different pilot plant samples (AO-FF and AOia). Two of these genomes were assembled from long-read sequencing, are HQ MAGs, and contain complete sequences of two rRNA operons, facilitating the analysis of this cluster with rRNA phylogeny. Copies of the *ppk1* gene found in these MAGs are phylogenetically differentiable from other *ppk1* sequences in the Accumulibacter lineage and create a separate clade (newly defined clade J) within Type II (Fig. 2). This is all strong evidence that allows us to propose the definition of a new species within the Accumulibacter lineage. For this UW20 cluster, we propose the epithet “jenkinsii” and a new species designation as “*Ca*. Accumulibacter jenkinsii” sp. nov., recognizing the many contributions of Dr. David Jenkins (aka Flocdoc) to our understanding of activated sludge microbiology (Table S2).

It is interesting that the *ppk1* gene sequence of “*Ca*. Accumulibacter jenkinsii” did not cluster together with other Accumulibacter *ppk1* sequences included in the large database that we used (Fig. S2). A possible explanation is that the samples analyzed in this study have the unique characteristic that they were collected from pilot plants operated under reduced aeration and with real wastewater. Thus, it is plausible that “*Ca*. A. jenkinsii” is adapted to treatment plant operations treating real wastewater at low-DO conditions. The reference genome of “*Ca*. A. meliphilus” (strain UW-LDO) was also obtained from a low-DO bioreactor, but in that case, the bioreactor was a lab-scale system receiving acetate as the only carbon and energy source (27). The reference genome for “*Ca*. A. necessarius” (strain UW12-POB) was recovered from a photosynthetic bioreactor in which acetate was also the main organic substrate and the DO was below 0.05 mg/L 70% of the time (26, 34).

Although all phylogenetic analyses support “*Ca*. A. jenkinsii” as a new species and new Type II clade within the Accumulibacter lineage, we acknowledge that additional *in situ* experiments are required to confirm the activity of “*Ca*. A. jenkinsii” as a canonical PAO using FISH and a method such as Raman microspectroscopy to detect the presence of storage polymers, as described in Petriglieri et al. (13).

### Phylogenetic classifications of UW18 and UW21 clusters

The *ppk1* and genome-based phylogenetic analyses placed the sequences in the UW18 cluster as belonging to clade II-C, along with “*Ca*. A. vicinus”, “*Ca*. A. cognatus”, and “*Ca*. A. contiguus” (Fig. 2). In agreement, the ribosomal protein-based phylogeny also places UW18 within clade II-C (Fig. 3). However, the UW18 MAG was assembled from short-read sequencing and does not include assembled rRNA operons or even partial sequences of these genes (Table 2). This and the fact that we only recovered one genome representing this cluster precludes us from proposing any new species designation for this cluster.

Likewise, the *ppk1*, genome-based, and ribosomal proteins-based phylogenies placed the sequences in the UW21 cluster as forming a separate cluster within clade II-F, differentiable from the other “*Ca*. Accumulibacter” species in this clade (“*Ca*. A. adjunctus”, “*Ca*. A. iunctus”, “*Ca*. A. similis”, and “*Ca*. A. conexus”). Thus, *ppk1* and metagenomic based phylogenies would support the placing of this cluster as a separate species within clade II-F, but unfortunately, the lack of 16S rRNA and 23S rRNA sequences for the most closely related species, “*Ca*. A. adjunctus” (94.7% ANI shared with UW21), limits further classification of this cluster at this time.

### Relative abundance of “*Ca*. Accumulibacter” species in pilot scale systems

To examine “*Ca*. Accumulibacter” diversity in the unconventionally aerated pilot-scale systems, we mapped metagenome reads to genomes representative of each “*Ca*. Accumulibacter” species. For “*Ca*. Accumulibacter” species for which we obtained MAGs in this study, we used those with the best quality (Table 2) as representative genomes for the species. For the other “*Ca*. Accumulibacter” species we used the genomes designated as the reference genome for each species. This analysis revealed “*Ca*. A. propinquus” and “*Ca*. A. necessarius” as the most prevalent species in the pilot plants, regardless of the configuration or mode of operation (Fig. 6). Combined, these two species accounted for greater than 40% of the Accumulibacter population. Other species were detected at moderate abundances, including “*Ca*. A. meliphilus”, “*Ca*. A. delftensis”, “*Ca*. A. jenkinsii”, and the UW18 and UW21 clusters among others (Fig. 6). Species for which the read mapping method did not support their presence in the pilot plants included “*Ca*. A. adiacens”, “*Ca*. A. aalborgensis”, “*Ca*. A. vicinus”, “*Ca*. A. cognatus”, “*Ca*. A. iunctus”, and “*Ca*. A. adjunctus”.

**Figure 6.**
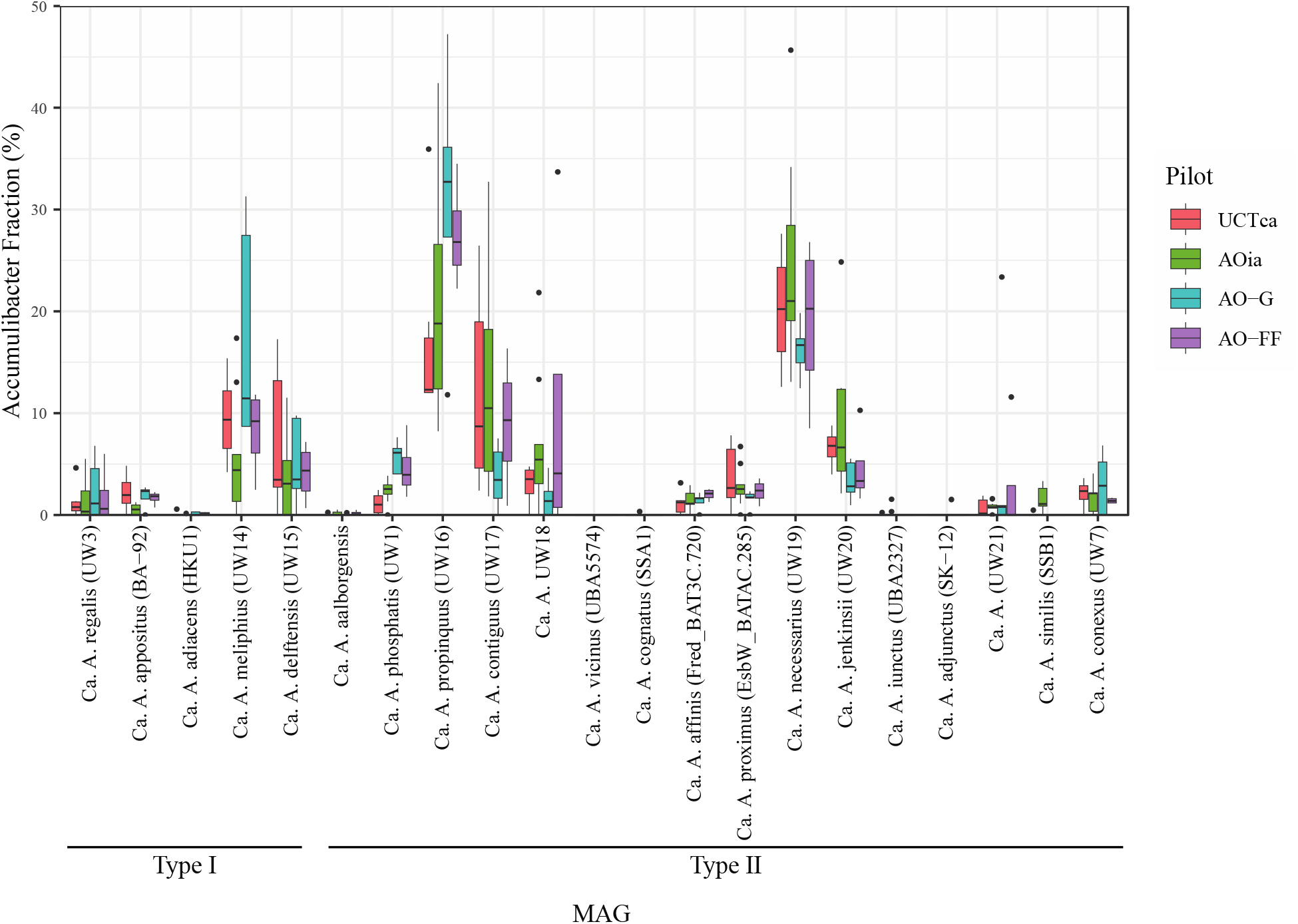
Box plot analysis of the relative abundance of each “*Ca*. Accumulibacter” species relative to the total “*Ca*. Accumulibacter” population. The pilot plants are color-coded, and each box indicates median and the first and third quartiles. The whiskers define the calculated minimum and maximum. Outliers outside the minima and maxima are indicated with black dots.

### Denitrification potential of the Accumulibacter population

The denitrification potential of the different “*Ca*. Accumulibacter” species is of particular interest in BNR processes operated with reduced aeration since oxidized forms of nitrogen could be used as electron acceptors in place of oxygen (11, 12) or potentially in addition to oxygen (27). A summary of denitrification genes found in high quality Accumulibacter MAGs was recently performed by Petriglieri et al. (13) and here we review the denitrification potential found in the assembled Accumulibacter MAGs and compare the observations to those of Petriglieri et al. (13), for “*Ca*. Accumulibacter” species for which MAGs with greater than 90% completion were recovered (Table 4).

**Table 4.**
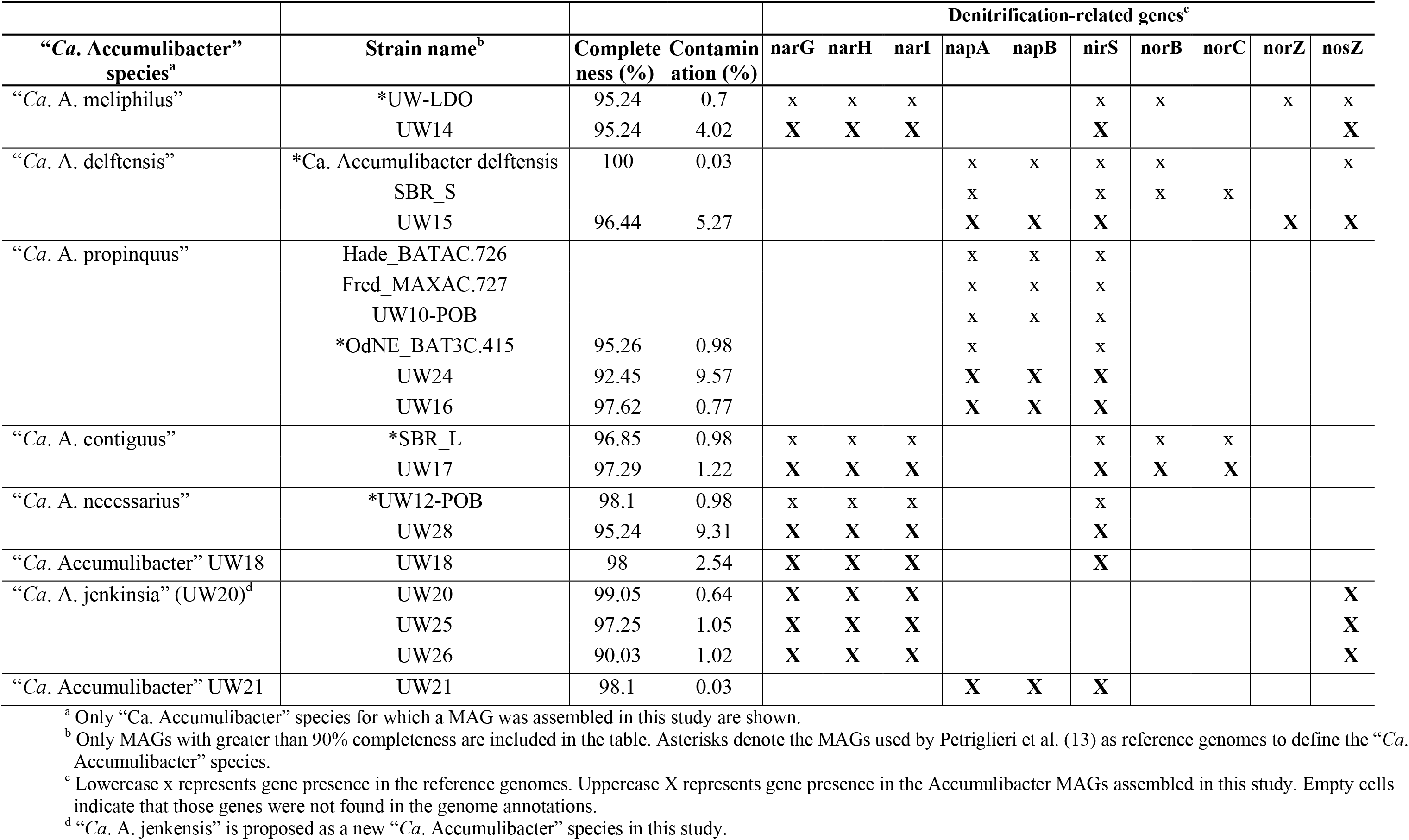
Analysis of denitrification potential of “*Ca*. Accumulibacter” species for which a MAG was assembled in this study.

Among the Type I Accumulibacter, “*Ca*. A. meliphilus” is the only species that has been shown to harbor the membrane-bound nitrate reductase genes (*narGHI*) necessary for respiratory NO_3_^-^ reduction to NO_2_^-^ (13, 27). The presence of these genes in “*Ca*. A. meliphilus” is confirmed by their detection in the UW14 MAG, which belongs to the same species (Table 4). Among Type II Accumulibacter, the assembled MAGs UW17 and UW28 confirm the presence of *narGHI* in “*Ca*. A. contiguus” (clade II-C) and “*Ca*. A. necessarius” (clade II-D), respectively. In addition, these genes were also found in the assembled MAGs for “*Ca*. A. jenkinsii” (new clade II-J) and for the UW18 cluster (clade II-C), but not for the UW21 cluster (clade II-F). These results are in agreement with *narGHI* being present in members of clade II-C, but not found in members of clade II-F (13).

The periplasmic nitrate reductase, *nap*, does not translocate protons, thus it ultimately does not contribute to energy generation via proton motive force, but can contribute to NO_3_^-^ reduction to NO_2_^-^. Through gene-flux analysis, it has been suggested that the *nap* system is core amongst all Type I Accumulibacter (35), however its presence is also widespread amongst Type II Accumulibacter genomes (13, 14). Also, in agreement with prior analyses, the periplasmic nitrate reductase genes (*napAB*) were confirmed to be present in “*Ca*. A. delftensis” (clade I-C) by the UW15 MAG and in “*Ca*. A. propinquus” (clade II-B) by the UW24 and UW16 MAGs (Table 4).

The presence of the nitrite reductase gene *nirS*, which catalyzes the reduction of NO_2_^-^ to nitric oxide (NO) (36) has been reported to be found in all “*Ca*. Accumulibacter” species, except for “*Ca*. A. cognatus” (13), one of the species in clade II-C for which we did not recover a MAG. The near universality of *nirS* in “*Ca*. Accumulibacter” species was confirmed by its presence in the MAGs associated with “*Ca*. A. meliphilus”, “*Ca*. A. delftensis”, “*Ca*. A. propinquus”, “*Ca*. A. contiguus”, and “*Ca*. A. necessarius”. However, *nirS* was notably absent in all the high-quality genomes that represent the newly proposed “*Ca*. A. jenkinsii” species (Table 4).

The reduction of NO to nitrous oxide (N_2_O) is catalyzed by nitric oxide reductases (NOR). NOR enzymes can be classified into two types: cNor or qNor (37). The cNor form is encoded by the *norBC* genes while the alternate form qNor is encoded by the *norZ* gene. The presence of both subunits of *norBC* has only been reported for the reference genome of “*Ca*. A. contiguus” (strain SBR_L; clade II-C) and a MAG classified as “*Ca*. A. delftensis” (strain SBR_S) (13). The presence of these genes in “*Ca*. A. contiguus” is supported by their annotation in the UW17 MAG, but the presence in “*Ca*. A. delftensis” is not supported by the UW15 MAG (Table 4). The *norZ* gene has been reported to be present in the reference genomes of “*Ca*. A. regalis” (clade I-A), “*Ca*. A. appositus” (clade I-B), “*Ca*. A. meliphilus” (clade I-C), and “*Ca*. A. phosphatis” (clade II-A) (27). Among the recovered MAGs, *norZ* was only annotated in the “*Ca*. A. delftensis” genome UW15.

N_2_O reduction to nitrogen gas (N_2_), the last reaction in the denitrification pathway, is encoded by the *nosZ* gene. The presence of this gene in the UW14 and UW15 genomes support prior detections in “*Ca*. A. meliphilus” and “*Ca*. A. delftensis”. This gene was also present in the three high-quality genomes recovered for the new “*Ca*. A. jenkinsii” clade (Table 4).

To date, a full respiratory denitrification pathway has only been identified in the reference genome of “*Ca*. A. meliphilus” (strain UW-LDO), consisting of *narGHI*, *nirS*, *norZ*, and *nosZ* genes. However, in the assembled “*Ca*. A. meliphilus” genome, UW15, neither *norBC* or *norZ* were annotated, and thus, our results do not support the earlier observations of a complete respiratory denitrification pathway in “*Ca*. A. meliphilus” (27).

## CONCLUSIONS

In this study, we used a set of newly assembled medium- to high-quality Accumulibacter MAGs to investigate the Accumulibacter population structure and dynamics in pilot-scale BNR systems operated with low-DO conditions. We also used this dataset to refine the phylogenetic analysis of the “*Ca*. Accumulibacter” species, taking advantage of the presence of complete rRNA operons in many of the newly recovered MAGs. We found that the Accumulibacter assemblages in these pilot-scale systems were heterogeneous, with metagenome-mapping based relative abundance analysis showing that “*Ca*. A. necessarius and “*Ca*. A. propinquus” accounted for greater than 40% of the “*Ca*. Accumulibacter” assemblage, and that members of “*Ca*. A. meliphilus”, “*Ca*. A. delftensis”, and “*Ca*. A. contiguus” were also present at moderate abundances. Moreover, half of the newly assembled MAGs could not be classified within the existing “*Ca*. Accumulibacter” species and were grouped into three phylogenetically different clusters that were also present at moderate abundances. In particular, one cluster among these also represented a new clade within Type II Accumulibacter, named here Type II-J. For this new clade we proposed the epithet “jenkinsii” and the designation “*Ca*. Accumulibacter jenkinsii” sp. nov. An examination of denitrification potential within the newly assembled Accumulibacter MAG was mainly confirmatory of earlier analyses of denitrification potential in “Ca. Accumulibacter” species, except for “*Ca*. A. meliphilus”, which was originally reported to have a complete set of denitrification genes (27). In the assembled “*Ca*. A. meliphilus” UW14 strain we did not find copies of the *norBC* or *norZ* genes needed for NO reduction to N_2_O. Our analysis did not provide evidence of niche differentiating roles related to the differences in operational conditions in the four pilot plants studied. Rather, we conclude that the diversity of “*Ca*. Accumulibacter” species seeing in the samples from the pilot plants operated with unconventionally low aeration indicates the widespread ability of “*Ca*. Accumulibacter” species to efficiently grow in conditions in which the aerated zone has low DO concentrations. This metagenome-based observation agrees with kinetics-based analysis that show that PAOs in BNR systems have high affinity for oxygen and are not negatively affected by changing operational conditions to low aeration (21, 27). The implication of these observations is that regardless of the mode of operation, efficient phosphorus removal can be effectively maintained when transitioning BNR processes to reduced aeration, a strategy that is gaining interest to reduce energy consumption in BNR processes.

## METHODS

### Operation of pilot-scale plants

Metagenomic samples from four instances of pilot-scale plant experiments were used in this study (Fig. 1). In all four experiments, the objective was to evaluate BNR performance when oxygen availability was reduced, compared to aeration conditions used in typical full-scale operation. In all cases, treatment trains were operated at the Nine Springs wastewater treatment plant (Madison, WI) and mimicked flow rate variations experienced in the full-scale process. Water temperature was not controlled and followed the same trends experienced in the full-scale process. All pilot plants treated primary effluent produced by the full-scale operation. In one set of experiments, two pilot plants were operated for 731 days. In a second set of experiments, two pilot plants were operated for 505 days.

A detailed explanation of the first 483 days of operation of the first set of experiments was described in Stewart et al. (19). Briefly, one pilot-scale system was configured as an anoxic-oxic (AO) process with intermittent aeration in the aerated zone (AOia), and the other was configured to resemble the full-scale treatment plant, which is a University of Cape Town-type (UCT) process (38) operated without internal nitrate recycle and with continuous aeration (UCTca). Aeration in each pilot-scale plant was controlled via ammonia-based aeration control. Intermittent aeration in the AOia pilot reactor was designed to maintain an ammonium (NH_4_^+^) concentration between 2 and 5 mgNH_4_^+^-N/L at the mid-point of the aerated section. Air-on mode was initiated once the NH_4_^+^ concentration reached the high setpoint of 5 mgNH_4_^+^-N/L. Conversely, once the low setpoint of 2 mgNH_4_^+^-N/L was reached, air-off mode began. During air-on mode the DO was maintained at or below 0.7 mg/L. Continuous aeration in the UCTca pilot reactor was controlled to maintain an ammonium concentration of 5 mgNH_4_^+^-N/L at the mid-point of the aerated section in the treatment train. In this setting, the aeration rate was adjusted as necessary to reach the NH_4_^+^ setpoint, but constrained to maintain DO between 0.1 and 0.6 mg/L. The solids retention time (SRT) of both plants was increased to overcome lower nitrification rates during the winter season when water temperatures reached as low as 12°C (19). In the second set of pilot-plant experiments, two pilot plants were configured as AO processes and operated to evaluate the effect of selective wasting on activated sludge settleability while transitioning a BNR system from conventional high-DO to low-DO treatment. A detailed explanation of the operation of these pilot-scale systems can be found in Amstadt (20). One pilot was equipped with a gravity-based selective wasting device, and therefore designated as the AO-G pilot-scale plant (Fig. 1). Selective wasting in the other pilot was implemented through a floc filtering device, and therefore designated as the AO-FF. Return activated sludge (RAS) fermentation was also implemented in the AO-FF pilot-scale plant by deviating 10% of the RAS to a fermentation tank. Stepwise reductions in DO were carried out in phases throughout the operation of these two pilot-scale plants.

### Metagenomic sequencing

Mixed liquor samples were collected from the four pilot plants at different times of operation and stored at -80°C until DNA extraction was performed. DNA extraction aimed at producing fragments adequate for long-read sequencing and followed either of two methods: 1) DNeasy PowerSoil Kit (Qiagen) without bead-beating and vortexing steps, or 2) a phenol-chloroform method (39) modified by removing the bead-beating step. DNA was quantified using a Qubit fluorometer (ThermoFisher Scientific) and assessed for quality using a Nanodrop 2000 spectrophotometer (ThermoFisher Scientific). For samples extracted using the DNeasy PowerSoil kit, an additional quality control analysis included DNA fragment sizing using a FemtoPulse system (Agilent). Extracted DNA was submitted for sequencing at the Joint Genome Institute (JGI). Depending on the quality of the DNA received by JGI, samples were prepared for either long-read sequencing on a PacBio Sequel IIe platform (Pacific Biosciences, Inc., Menlo Park, CA, USA) or short-read sequencing on an Illumina NovaSeq 6000 platform. For long-read sequencing, libraries were prepared by shearing genomic DNA to either 3 kb or 6-10 kb, depending on DNA quality, and performing ligations using the SMRTbell Express template prep kit (Pacific Biosciences). Size selection was performed with BluePippin (Sage Sciences). Library quality was assessed using the FemptoPulse system and quantified using the Qubit fluorometer. Finally, the libraries were sequenced on a Sequel IIe using Sequel Polymerase Binding Kit 2.2. When DNA quality was deemed not to be adequate for long-read sequencing, genomic DNA was sheared to 400 bp using the Covaris LE220 instrument and size selected with SPRI using TotalPure NGS beads (Omega Bio-tek). Library prep included fragment end treatment, A-tailing, and ligation of Illumina-compatible adapters (IDT, Inc.) using the Kapa-HyperPrep kit (Kapa Biosystems). DNA was then sequenced 2 x 150 bp on the Illumina NovaSeq 6000 platform using S4 flow cells. Upon completion, we obtained 15 metagenomes from 6-10kb PacBio, 7 from 3kb PacBio, and 2 from Illumina libraries.

### Metagenome assembly, binning, and annotation

Metagenome assembly, binning, and annotation was performed, individually for each metagenome, using default parameters for each program. PacBio reads were filtered using BBtools (v38.87/38.88) and assembled using metaFlye (v2.9) (40). PacBio assemblies were polished with racon (v1.4.13) (41). Cleaned PacBio reads were then mapped on to the assemblies using minimap2 (42). Illumina reads were processed following JGI’s published pipeline (43) as follows. Raw reads were filtered and corrected using bbcms (v38.90), assembled using metaSPAdes (v3.15.2) (44), and mapped to the final assembly using BBMap (v38.86). All assembled contigs generated from both PacBio and Illuminia sequencing technology were binned using metaBAT2 (45). The resulting metagenome-assembled genomes (MAGs) were annotated following procedures in the JGI metagenome processing pipeline (43).

### Phylogenetic analyses and relative abundance

To determine the phylogenetic classification of the Accumulibacter MAGs, we clustered them along with the reference genomes of recognized “*Ca*. Accumulibacter” species using dRep (v3.4.0) (46) with a 95% average nucleotide identity (ANI) threshold (-sa 0.95 -comp 50 -con 10), which first divides the genome set into primary clusters using Mash (v2.3) (47). In addition, pairwise genome-wide ANI calculations between MAGs were made with FastANI (v1.33) (48).

Genome-level phylogenetic analyses were conducted using 120 single-copy marker gene protein sequences identified and aligned with GTDB-Tk using default parameters (v1.6.0, database release 202)(49).

A phylogenetic analysis based on a fragment of the *ppk1* gene was conducted using a database of reference and clone sequences compiled by McDaniel et al. (25), supplemented with additional *ppk1* gene sequences from Petriglieri et al. (13) and the *ppk1* gene sequences from the Accumulibacter MAGs assembled in this study. In addition, a phylogenetic analysis based on the full sequence of the *ppk1* gene was performed using the *ppk1* genes from the Accumulibacter MAGs assembled in this study and the *ppk1* genes from the MAGs that are representative of each “*Ca*. Accumulibacter” species. In both cases, the sequences were aligned using MAFFT “-- auto” (v7.222) (50).

Phylogenetic analyses based on the 16S rRNA and 23S rRNA genes were performed by aligning the sequences of these genes extracted from the Accumulibacter MAGs, from the reference genomes of the “*Ca*. Accumulibacter” species, and from other Accumulibacter-related contigs in the metagenome assemblies that included complete rRNA operons. To provide context, 16S rRNA sequences assigned to the Accumulibacter lineage in the MiDAS database (v4.8.1) (51) were also used in the 16S rRNA analysis. Geneious Prime version 2022.0.2 (https://www.geneious.com) was used for the extraction of rRNA gene sequences, the alignment of sequences with MAFFT (50).

A ribosomal phylogenetic tree of Accumulibacter MAGs was constructed with the metabolisHMM package (v2.21) (29), which uses a collection of 16 bacterial ribosomal proteins (52) and requires that each MAG contain at least 12 ribosomal proteins. The package searches for each protein using specific HMM profiles, aligns them using MAFFT (50), and concatenates all markers together.

For all phylogenetic analyses, maximum likelihood trees were created from the alignments with RAxML-NG (“--model LG+G8+F” model) (v0.9.0) (53) and visualized with iTOL (54).

We used the metagenome reads to estimate relative abundance of “*Ca*. Accumulibacter” species in the different samples. For “*Ca*. Accumulibacter” species for which a MAG was recovered in any of the samples, we selected the best MAG (as determined by dRep scores) as the representative for that species. For “*Ca*. Accumulibacter” species for which a MAG was not recovered from the pilot plant samples, we selected the reference genome as the representative genome. Since the metagenome libraries contained long PacBio and short Illumina reads, the filtered PacBio long-reads of both 3 kb and 6-10 kb libraries were divided into smaller 500 bp long sequences using BBMap/shred.sh (v38.32) with no overlap between successive reads (55). The shortened PacBio reads and the cleaned short-reads from Illumina libraries were competitively mapped (i.e., reads mapped onto multiple genomes simultaneously) onto the set of representative “*Ca*. Accumulibacter” MAGs using Bowtie2 (v2.3.5.1) with the default “--end-to-end” and “--fast” options (56). The program coverM (v0.6.1) (https://github.com/wwood/CoverM) was used to obtain the relative abundance of reads mapped onto each MAG with the “coverm genome” command.

### Data availability

The metagenomes used in this study are available at the JGI gold database under Study ID: Gs0156633. Table S1 includes links to the specific projects within this study that contain the metagenomes from each pilot plant, and the corresponding JGI metagenome analysis project ID for each metagenome. Table 2 includes links to the ID numbers in the Integrated Microbial Genomes (IMG) database for each MAG described in this study. The datasets can also be found on the National Center for Biotechnology Information (NCBI) website under BioProject accession number PRJNA1037153.

## ACKNOWLEDGMENTS

This work was partially supported by funding from the National Science Foundation (CBET-1803055), the Madison Metropolitan Sewerage District (Madison, WI), and the Great Lakes Bioenergy Research Center (award number DE-SC0018409; U.S. Department of Energy, Office of Science). The work (proposal 10.46936/10.25585/60001333) conducted by the U.S. DOE JGI (https://ror.org/04xm1d337), a DOE Office of Science User Facility, is supported by the Office of Science of the U.S. DOE operated under contract DE-AC02-05CH11231. R.D.S. was partially funded by National Science Foundation (HRD-1810916) and the Graduate Engineering Research Scholars (GERS) program of the College of Engineering at University of Wisconsin – Madison. The authors utilized the University of Wisconsin – Madison Biotechnology Center’s DNA Sequencing Facility (Research Resource Identifier – RRID:SCR_017759) to perform quality control of high-molecular weight (HMW) DNA. We thank James Alvin, Rania Bashar, Kailey DeVault, Maddie Douglas, Brennan Ellis, Ellen Feigl, Lillian Glackin, Morgan Keck, Michael Liu, Lucas Lobreglio, Sara Neufcourt, Anthony Radicia, Gustavo Uribe-Santos, Claire Steins, and Trenton Weiss for support with pilot plant operation, and Maria Chuvochina for help with the etymology of the proposed names.

